# Visualization of spatial gene expression in plants by modified RNAscope fluorescent *in situ* hybridization

**DOI:** 10.1101/2020.04.01.020727

**Authors:** Shyam Solanki, Gazala Ameen, Jin Zhao, Jordan Flaten, Pawel Borowicz, Robert Brueggeman

## Abstract

*In situ* analysis of biomarkers such as DNA, RNA and proteins are important for research and diagnostic purposes. At the RNA level, plant gene expression studies rely on qPCR, RNAseq and probe-based *<u>i</u>n <u>s</u>itu* <u>h</u>ybridization (ISH). However, for ISH experiments poor stability of RNA and RNA based probes commonly results in poor detection or poor reproducibility. Recently, the development and availability of the RNAscope RNA-ISH method addressed these problems by novel signal amplification and background suppression. This method is capable of simultaneous detection of multiple target RNAs down to the single molecule level in individual cells, allowing researchers to study spatio-temporal patterning of gene expression. However, this method has not been optimized thus poorly utilized for plant specific gene expression studies which would allow for fluorescent multiplex detection. Here we provide a step-by-step method for sample collection and pretreatment optimization to perform the RNAscope assay in the leaf tissues of model monocot plant barley. We have shown the ubiquitous *HvGAPH* and predominantly stomatal guard cell expressed *Rpg1* expression pattern in barley leaf sections and described the improve RNAcope methodology suitable for plant tissues using confocal laser microscope. By addressing the problems in the sample collection and incorporating additional sample backing steps we have significantly reduced the section detachment and experiment failure problems. Further, by reducing the time of protease treatment, we minimized the sample disintegration due to over digestion of barley tissues. Thus, we optimized the RNAscope detection method in plants to visualize the spatial expression and semi-quantification of target RNAs which can be employed in other plants such as the widely utilized model dicot plant Arabidopsis.

## Introduction

*In vivo* fluorescent RNA *<u>i</u>n <u>s</u>itu* <u>h</u>ybridization (ISH) is a powerful technique utilized to detect the mRNA molecules [1, 2] in cells or tissue, enabling researchers to evaluate spatio-temporal gene expression. ISH approaches are popularly utilized in many cytological contexts and are particularly helpful to understand the expression pattern of morphological [3] or stress responsive genes [4] and small RNAs such as lncRNAs or microRNAs [5], that function to regulate various cellular responses. However traditional ISH methods are time consuming, labor intensive, and largely affected by poor stability of RNA in the sample [6, 7]. Consequently, ISH experiments majorly results in suboptimal detection of RNA in performed assay. Furthermore, high autofluorescence from the plant tissues further exacerbate the problems in the detection of low abundance RNA species. However, the recently described RNAscope ISH (ACDBio^™^) method overcomes the traditional drawbacks and provides the capability to multiplex targets in a rapid and highly sensitive assay. RNAscope detection relies on signal amplification and suppression of background noise, a major improvement over traditional RNA ISH method. The unique RNAscope ‘Z’ oligonucleotide RNA probes consist of ∼18-25 bases complimentary to the target sequence linked to a preamplifier binding motif (Fig. 1). A combination of ∼18-25 ZZ pairs designed to bind in a region spanning ∼1000 bases of target RNA, allows for specific detection and signal amplification from each detected target RNA molecule. Pre-Amplifier (PreAMP) binds to the preamplifier binding site created by each ZZ pairing, allowing the subsequent binding of multiple amplifiers (AMP). Fluorescence probes bind to each of these amplifiers, enabling the fluorescent visualization of mRNA. This chain of binding events greatly enhances the intensity of fluorescence signal from the target, enabling the detection of a single RNA molecule in a cell (Fig. 1). By using the probe specific amplifiers and sequence labelling, this assay can be multiplexed, thus provides the opportunity to visualize multiple target RNA species simultaneously. With the single cell sequencing approach gaining popularity to map the spatial expression of RNA molecules and identification of cell specific markers, multiplex RNAscope assay provides an excellent approach for validation of sequencing data and to visualize candidate gene expression in the specific cell types. Additionally, the capability of simultaneous immune-histo-chemistry and gene encoded protein detection followed by RNA visualization provides an option for evaluating RNA expression and efficiency of protein turnover validation in the same cell.

**Figure 1.**
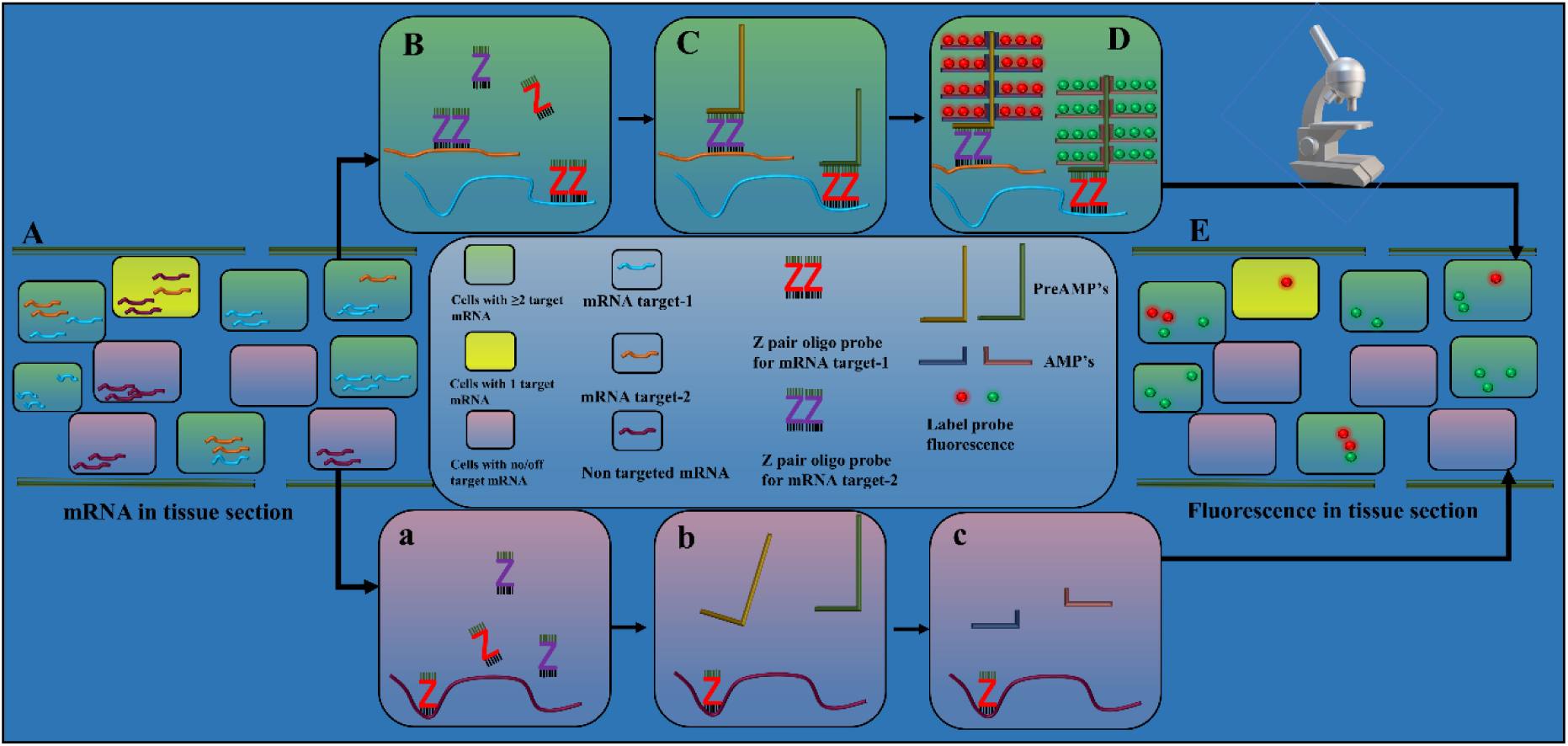
A schematic plan describing the multiplex RNAscope-ISH assay to visualize the spatio-temoral expression of more than one RNA species in a tissues section. Fixed plant tissue section with preserved RNA in each cell (A). RNA species can be targeted or non-targeted. Binding of target ZZ probe pair (B), PreAMPs (C), AMPs and fluorescence probes (D) in a cell containing the target RNA species. Inability of ZZ probe pair, PreAMP, AMP and fluorophore binding (a, b, c) in cells containing non-targeted RNA species only. Visualization of probe fluorescence in the tissue section in spatio-temporal manner for targeted RNA species (E).

Multiplexed fluorescent RNAsope has been widely adopted in mammalian research studies, however utilization in plants has been hindered due to the lack of a suitable plant tissue specific protocol describing the sample preparation, downstream staining and image processing. The standard RNAscope procedure described for the ACDBio^™^ [8, 9] system is not adapted for plant tissues and result in issues such as suboptimal plant tissue attachment to the charged slides and fluorescence detection. Thus, for utilization in plants considerable modification were required for better attachment of tissue sections, RNA integrity preservation, probe binding and sample visualization. Previously single/2-plex capable chromogenic RNAscope *in situ* hybridization methods were described using <u>f</u>ormaldehyde <u>f</u>ixed and <u>p</u>araffin <u>e</u>mbedded (FFPE) sample preparation [10] methods to detect plant leaf mRNAs in maize [11], cassava brown streak virus infecting cassava and nicotiana stems [12], however, the approach of utilizing RNAscope fluorescent probes capable of multiplexing targets have not been reported in plants. Here we optimized and described RNAscope multiplex fluorescent detection in fixed frozen barley leaf tissue. Barley, a model cereal crop plant, has been extensively studied and has available high-quality annotated genome sequences available [13, 14]. Many gene expression have been conducted with barley during biotic and abiotic stress responses. The barley *Rpg1* gene, previously shown to provide broad spectrum resistance against stem rust pathogens [15] was shown to have three fold higher expression in the epidermal fraction compared to the whole leaf from the resistant barley line Morex [16] normalized to the *HvGAPDH* housekeeping gene. In the present study the expression of the barley genes *HvGAPDH* and *Rpg1* were visualized in leaf tissue sections of barley line Q21861 by optimizing the multiplex RNAscope fluorescent assay. Here we are reporting an optimized three-day fluorescent RNAscope protocol on plants from sample collection to RNA visualization, a tool for convenient and efficient detection of RNA in fresh frozen plant leaf tissue sections.

## Material and methods

### Reagents and equipment

RNAscope Probes (Here we reported custom synthesized *HvGAPDH* and *Rpg1* specific probes in barley, ACDBio^™^)

RNAscope V2 kit (ACDBio^™^)

Immedge® Hydrophobic Barrier Pen (Vector Laboratories Cat. No. 310018)

Superfrost plus slides (ThermoFisher Scientific, 4951PLUS4)

Tissue-Tek® manual slide staining set (Sakura, 4451)

Tissue-Tek® 24-slide slide holder with detachable handle (Sakura, 4465)

HybEZ(tm) hybridization system and EZ-batch^™^ slide processing system (ACDBio^™^)

Humidifying paper (ACDBio^™^, 310025)

Incubator (VWR, 10055-006)

Water Bath (ThermoFisher Scientific, TSGP02)

Thermometer (VWR)

Zeiss LSM-700 confocal microscope

Fresh 10% NBF (Millipore Sigma, HT501128-4L)

Ethanol (Millipore Sigma, E7023)

1x PBS, Ph 7.4 (ThermoFisher Scientific, 10010023)

Everbright mounting media (Biotium, 23003)

Peel-A-Way embedding cryomold (Millipore Sigma, E6032-1CS)

Tissue-Tek® optimal cutting temperature (OCT) compound (Sakura 4583)

### Plant growth and sample collection

In controlled environment growth chambers (16 hour light : 8 hour dark; light intensity at 300 µmol s^-1^ m^-2^ in Conviron growth chamber) barley line Q21861 seeds were planted in cones filled with a peat moss:perlite (3:1 v/v) potting mix (#1 Sunshine Mix, Fisons, Vancouver, Canada). Barley line Q21861 contains the *Rpg1* gene. Plants were grown for ten days or until the primary leaf was fully expanded. Before sample collection a plastic Peel-A-Way embedding cryomold (Millipore Sigma) was filled with frozen <u>o</u>ptimal <u>c</u>utting <u>t</u>emperature (OCT) compound and kept aseptically on dry ice for two-three minutes to achieve a semi frozen state (Fig. 2A). This step is important for leaf sample placement and to achieve desired orientation. Leaf samples tends to shift its position in a liquid OCT compound, thus maintaining the semi frozen state is critical. For sample collection, 3-4 cm long leaf sections were cut aseptically and placed in the desired orientation in the semi frozen OCT compound (Fig. 2B). We placed samples to obtain a cross section cut of leaf samples. To obtain transfer of multiple tissue sections probing for each target gene in one slide, we have collected 2-3 leaf bits from four individual plants in one cryomold in semifrozen OCT and then let OCT completely frozen. Having multiple tissue sections in one cryomold serves the purpose of sample replication (Biological) and increase the chances of obtaining multiple images using less quantity of expensive reagents. Cryomold blocks were stored at −80°C ultra-freezer until sample sectioning. Samples stored for up to six-months have been used without showing compromised sample integrity.

**Figure 2.**
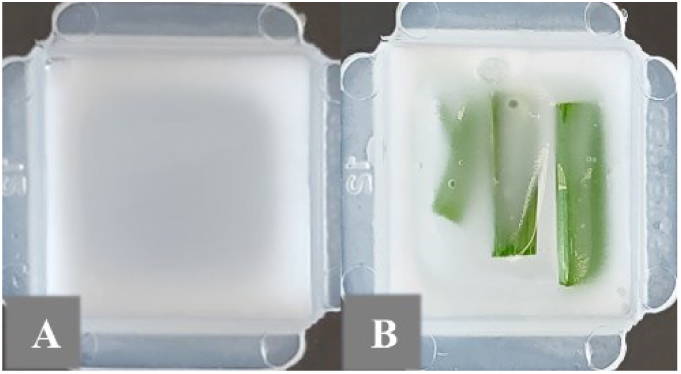
Collection of fresh frozen sample in OCT compound filled cryomolds on dry ice. (A) Semi frozen OCT compound helps maintain desired leaf orientation for (B) subsequent microdissection.

### Sample sectioning and fixation

Before sectioning, frozen OCT containing leaf samples were equilibrated at −20°C in a microtome cryostat (Leica) for one hour. The cryomold was peeled out of the OCT compound block and 10-15 µm thick sections were cut and mounted on the Superfrost plus (ThermoFisher Scientific) microscope slides. Slides were kept in the cryostat at −20°C for 30 minutes to let them dry and devoid of moisture. These sections can be immediately processed or stored in −80°C for up to 2-3 months in an airtight container. However, it is preferred to fix the samples immediately after sectioning. Samples were fixed according to the RNAscope kit manufacturer (ACDBio) standard protocol with slight modifications. In brief, slides containing tissue sections were placed in the slide rack and immediately dipped in the pre-chilled fresh 10% <u>n</u>eutral <u>b</u>uffered <u>f</u>ormalin (NBF) solution (Millipore Sigma) for one hour at 4°C in a staining dish. A longer fixation is required for the plant samples compare to the standard recommend 15 minutes fixation time for animal tissue sections to achieve optimal RNA integrity. After fixation, slides were briefly rinsed with 1X <u>p</u>hosphate <u>b</u>uffered <u>s</u>aline (PBS) two times to wash away the fixative. Tissue sections were then dehydrated in a series of ethanol solution 50%, 70% and 100% for five minutes at room temperature. After ethanol washes the slides were transferred into a fresh 100% ethanol solution for five minutes before creating the hydrophobic barriers. At this point slides can also be stored in a 100% ethanol solution for up to one week. In our experiments we only stored the slides overnight before pretreatment processing.

### Sample pretreatment

The following day slides were removed from the 100% ethanol and baked at 37°C for 30 minutes to one hour. This step improves the sample attachment to the slides and avoids tissue removal from the slides during subsequent washing steps. After baking the slides to create a hydrophobic barrier around the sample a Immedge hydrophobic barrier pen (Vector Laboratories) was used while avoiding any touching or disruption of the tissue section. Utilizing the multiplex fluorescence V_2_ RNAscope kit (ACDBio) the hydrogen peroxide and Protease IV was applied according to the manufacturer instructions with a slight modification. Because it was observed that a 30-minute Protease IV treatment caused over-digestion of the leaf sections, the treatment was reduced to 15 minutes.

### Probe hybridization and RNAscope assay

The subsequent RNAscope assay was composed by multiple steps of hybridization (probe and AMP binding) and signal development (HRP channel binding), each followed by washing the slides in 1x wash buffer (Fig. 3) as described in the manufacturer protocol (Multiplex fluorescence V_2_ kit, ACDBio). The barley *HvGAPDH* RNAscope probes were custom developed in the C1 channel and the barley *Rpg1* probes in C2 channel by the ACDBio custom probe development unit utilizing the barley line Q21861 cDNA sequence of each gene. The *HvGAPDH* and *Rpg1* probes (200 µl) were applied to the Q21861 tissue sections encircled by a hydrophobic boundary, ensuring that the probe solution covered the entire sectioned area. A bacterial DapB 3-plex negative control probe was hybridized with sample tissues on a separate slide. Single probe and multiplexed assay were used to validate the efficiency of the assays. The experiment consisted of four hybridization assays containing; 1) the *HvGAPDH* probe, 2) the *Rpg1* probe, 3) *HvGAPDH* and *Rpg1* probes, and 4) the 3-plex negative control probe. The hybridizations were performed and analyzed using slides containing multiple independent tissue sections. For probe hybridization the samples were incubated in the HybEz oven for 3-4 hours at 40°C. In our experience, extended probe hybridization times of up to 4 hours with the plant samples improve the probe hybridization and subsequent visualization. However, a two-hour incubation period was also shown to be adequate once the assay was validated and standardized. After the 4-hour incubation and washing procedure the slides were stored in 5x SSC buffer, an optional pause point as suggestion by manufacturer. We found that breaking the assay in three days protocol allows to carry it very conveniently. The following day slides were washed in 1x wash buffer and AMP signal amplification and HRP labeled probe hybridization steps carried out according to the manufacturer protocol (Multiplex fluorescence V_2_ kit, ACDBio). We have used TSA plus® fluorescein (green) and TSA plus® Cyanine 3 (orange) fluorophores (PerkinElmer) diluted at 1:1000 to visualize *GAPDH-C1* signal in green and *Rpg1-C2* signal in orange, however any of the other described TSA plus® i.e. TSA plus® fluorescein (green), TSA plus® Cyanine3 (Orange) or TSA plus® Cyanine 5 (Red) can be used interchangeably. Developed sample slides after the AMP staining were treated with nuclear stain DAPI. DAPI was removed by flicking the slides withing five seconds and a cover slip was placed utilizing the Everbright hard set mounting media. These slides were stored overnight in the dark before visualization for complete drying up of hard-set mounting media.

**Figure 3.**
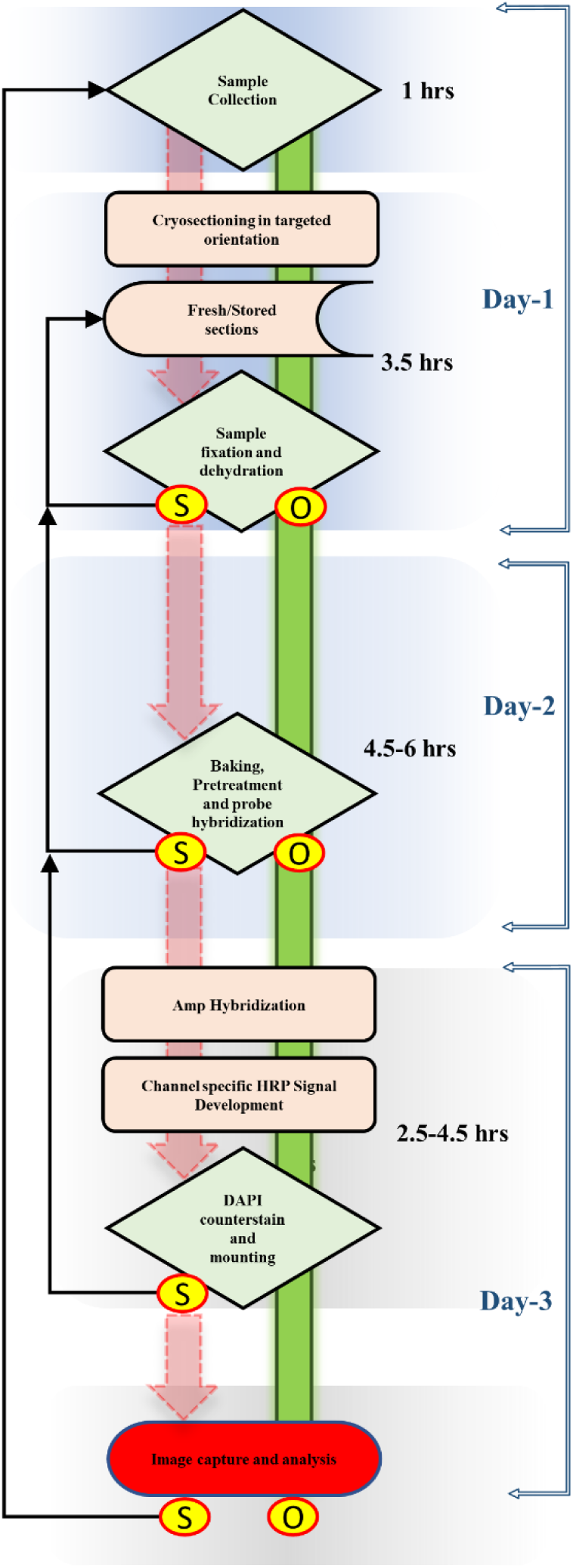
Schematic decision-making flow chart to efficiently carry out the various steps in the RNAscope assay. A rhombus represents the decision-making step, smooth edge rectangles represent a process, a storage symbol depicts the step for long term sample storage and terminator represents the end of a successful RNAscope experimental pipeline. Green bars represent a successful progression of the assay, whereas a red arrow represents a possible failure point that may require returning to the previous step (black arrows). O - Optimal, S - Suboptimal

### Image capturing

The RNAscope sample slides were visualized on a LSM 700 laser scanning confocal microscope using a Plan-Apochromat 20x/0.8 lens (air) and 10x objective lens with four different channels (Zeiss Thornwood, NY). Three fluorescent channels were used simultaneously; 1) the green channel 488/520 was assigned for TSA plus® fluorescein fluorophore (*HvGAPDH* probe), 2) the blue channel 360/460 was used for DAPI nuclear stain, and 3) the orange channel 550/570 was assigned for TSA plus® Cyanine3 (*Rpg1* probe). T-PMT was assigned for <u>d</u>ifferential <u>i</u>nterface <u>c</u>ontrast (DIC) imaging. For multilayer visualization Z-stack images were taken and computation from Z stack was performed in Imaris 9.0.1 software (Bitplane, South Windsor, CT) as described previously [17]. During image capture the negative control was assessed first to set the strength of LASER and exposer time. The process from sample collection to image analysis covered a three-day time frame without compromising the integrity of experimental samples (Fig. 3).

## Result and discussion

We have optimized the RNAscope multiplex fluorescent detection -V_2_ method for fixed frozen barley leaf tissue sections to visualize the expression profile of *HvGAPDH* and *Rpg1*. Fixed frozen plant samples provide an excellent alternative to FFPE samples which is a method of choice largely for the long-term preservation of animal and human tissue samples. RNA stability and tissue section integrity are critical factors required for the success of any ISH experiment including RNAscope. Due to sensitive detection level provided by the 18-25 adjacent pairs of Z probes, the RNAscope V_2_ assay provides a higher sensitivity assay than traditional ISH methods. However, the previous lack of sample preparation adaptability for plant-based assays that resulted in tissue detachment from slides and disintegration during multiple reagent washing steps caused loss of the samples in the V_2_ assay resulting in poor detection in our initial assay runs. A colorimetric RNAscope *in situ* hybridization method previously described using formaldehyde fixed and paraffin embedded (FFPE) sample preparation method targeting cassava brown streak virus infecting cassava and nicotiana stems [12] did not work for the fluorometric assay indicating the importance of optimizating the method and sample preparation. The fluorescent RNAscope assay has the advantage of visualizing multiple RNA species in one sample with the capability of localizing the site of RNA expression for spatial-transcriptomics analysis which complements gene expression studies [18]. In our experiment we have utilizes the barley *HvGAPDH* and *Rpg1* genes to visualize their expression in leaf sections. *HvGAPDH* is universally adopted as a housekeeping gene and widely utilized for the qPCR assays in barley and its related species [19] irrespective of recent reports of expression stability during stress responses [20]. Barley plants were grown without any deliberate stress response to maintain the normal expression level of candidate genes visualized by the RNAscope assay. In our initial attempts we encounter the problem of washing off of the samples during the assays resulting in experiment failures. To alleviate this issue, we included the sample baking step after fixing the tissue which resulted in better retention, which is now adopted in current modified RNAscope V2 assay manual. For the majority of samples tissue sections that were retained for final visualization, we observed another issue which was over digestion of fixed tissues which made it difficult to determine cellular morphology (Fig. 4A, 4B). To overcome this issue the protease treatment step was modified. These issues that were addressed by optimizing by sample baking and modifying the protease treatment steps, were reported to the ACDBio company in hopes that they will modify their standard protocol for plant tissue. We visualized the expression of *HvGAPDH* and *Rpg1* in both singleplex (Fig. 4C, 4D, 4E, 4F) and multiplexed assays (Fig. 5A, 5B), however the intensity of detection varied in different samples indicating a possible variability in the RNA integrity preservation during the sample preparation.

**Figure 4.**
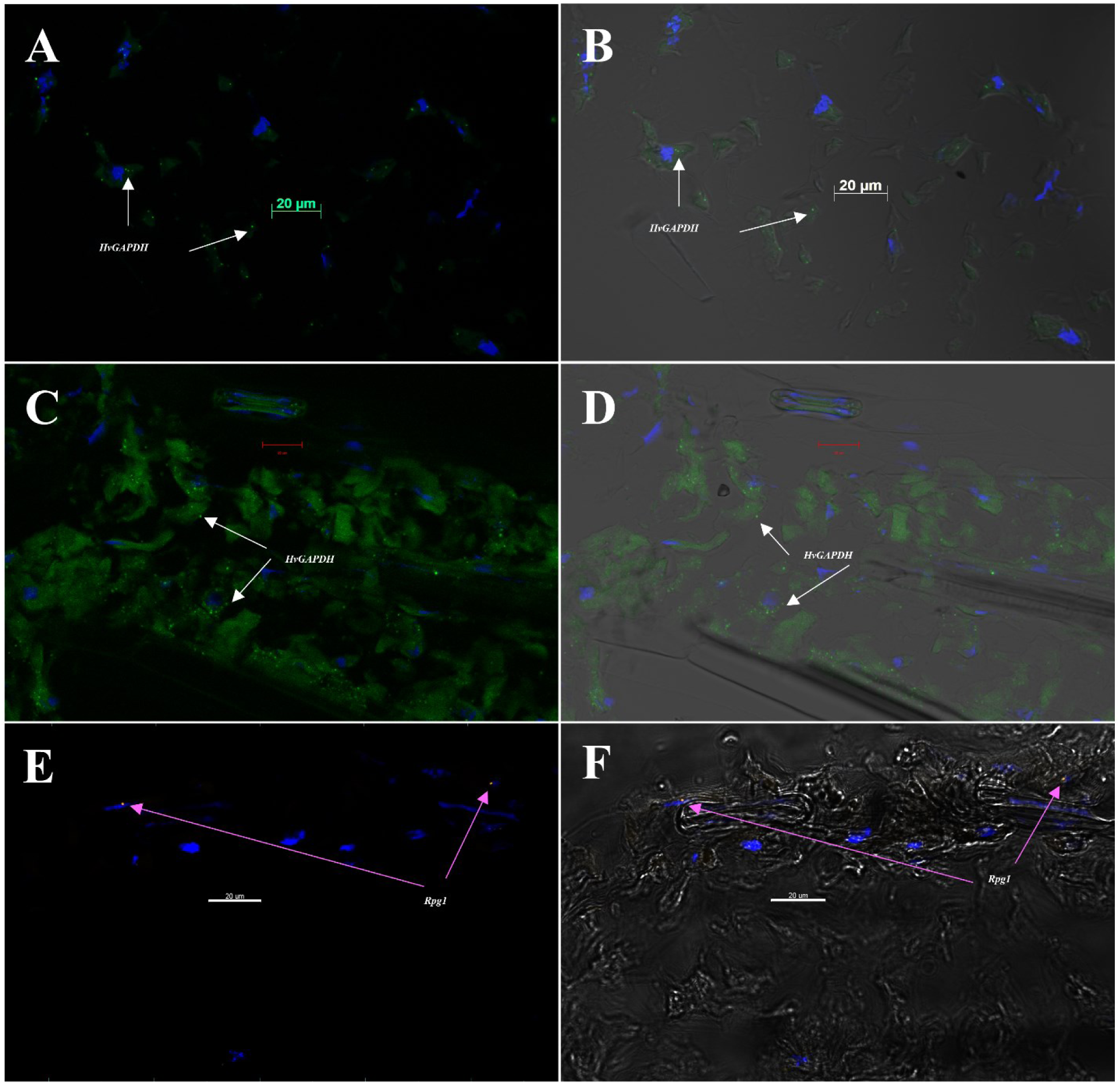
Single-plex detection in RNAscope V_2_ assay for *HvGAPDH* and *Rga1* mRNA on barley line Q21861 leaf sections during assay optimization. Confocal Image capturing on Zeiss LSM-700 without differential interference contrast (DIC) (A, C, E) and with DIC (B, D, F) transmitted light detcector T-PMT. (A, B.) Suboptimal digestion of barely leaf sections using a 30-minute Protease IV treatment resulting in the fragmentation and detachment of cells during multiple washing steps at below accepted levels. *HvGAPDH* expression is visualized as green distinctive dots (C1 channel hybridized with TSA plus® fluorescein) in the majority of barley cells and DAPI (blue) is used as nuclear stain. (C, D.) An optimized initial one-hour baking after sample fixation and 15 minutes Protease IV treatment resulted in the improved tissue integrity and mRNA associate fluorescein probe detection for *HvGAPDH*. (E, F.) A very low expression of *Rpg1* was visualized as orange fluorescence dots (C2 channel hybridized with TSA plus® Cyanine3) associated with the stomata associated subsidiary cells.

**Figure 5.**
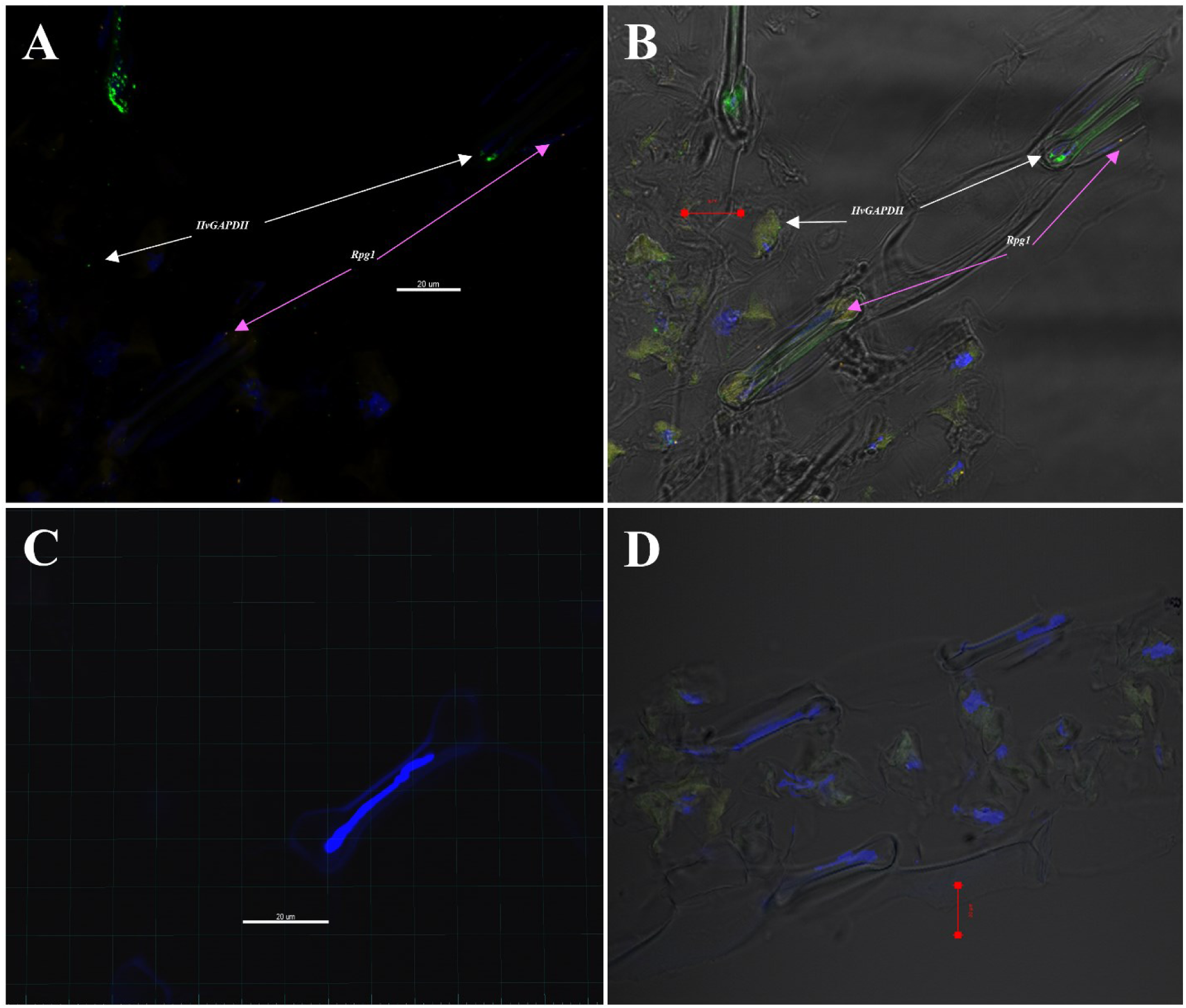
Multiplex detection in RNAscope V_2_ assay for simultaneous visualization of *HvGAPDH* and *Rga1* mRNA on barley line Q21861 leaf sections. Confocal Image capturing on Zeiss LSM-700 without DIC (A, C) and with DIC (B, D) transmitted light detcector T-PMT. (A, B.) Simultaneous visualization of *HvGAPDH* (C1 channel probe hybridized with TSA plus® fluorescein, green fluorescence dots) and *Rpg1* (C2 channel probe hybridized with TSA plus® Cyanine3, orange fluorescence dots). Majority of *Rpg1* expression was found to be associated with subsidiary cells associated with barley guard cells in the epidermal layer. DAPI stain was used for nuclear staining. (C, D.) A 3plex negative control with DAPI staining, without any probe specific fluorescence.

In successful experimental runs we have detected the uniform expression of *HvGAPDH* visualized as 2-8 distinctive dots in the leaf tissue section. Interestingly the majority of low level of *Rpg1* expression was predominantly associated with the barley epidermal subsidiary cells adjacent to the stomatal guard cells (Fig. 4E, 4F, 5A, 5B). The stem rust resistance gene *Rpg1* [21] was previously shown to have higher expression in the epidermal cells [16] and its rapid protein phosphorylation is critical to provide resistance response [22], thus our report of its predominant expression in epidermal stomata associated cells is in conjunction with the previous findings. Negative controls were largely without the presence of any dot-like signals (Fig 5C, 5D).

It important to interpret the results during the sample visualization to determine the background from the actual signal. Only distinctive dots should be considered as the specific signals and for a highly expressed targets more than one dots should be present in each intact cell. As described in the product literature and previously published human tissue specific protocols number of distinctive dot corresponding to the RNA copy number successfully bound with the probes, whereas intensity of each dot signifies the number of probe pairs bound to each RNA molecule. This quantification can vary depending upon the efficiency of probe hybridization during the sample preparation and level of gene expression, thus represents a semiquantitative nature of method and intuitive in nature.

## Conclusions

We report an optimized procedure for utilizing the novel RNAscope fluorescent multiplexing approach in plants for RNA expression visualization, which is otherwise only optimized for animal tissue types. Specifically, we have optimized the sample preparation and tissue pretreatment steps, a key component before probe binding and sample visualization using the model plant barley. These modifications minimize the tissue fragility, avoids fixation issues to maintain RNA integrity and detachment of plant material during the microtome sectioning and subsequent wash steps. Thus, ensuring a properly processed samples for microscopic visualization. Using the optimized protocol, we have shown the ubiquitous expression of barley housekeeping gene *HvGAPDH* along with expression of *Rpg1* mRNA which is predominantly visualized in the subsidiary cells adjacent to stomata present in the epidermal layers.

## Availability of data and materials

## Abbreviations

(ISH): *in situ* hybridization
(OCT): Optimal Cutting Temperature
(PBS): Phosphate Buffered Saline
(DIC): Differential interface contrast
(NBF): Neutral Buffered Formalin
(FFPE): Formaldehyde Fixed And Paraffin Embedded

## Declarations

### Ethics approval and consent to participate

Research work was carried out as per the local legislation.

### Consent for publication

Yes

### Availability of data and materials

All the seed material and raw imaging data can be obtained from the authors upon request.

## Author contribution

RSB and SS conceived the research idea and designed the experiment. SS and GA maintained greenhouse and collected samples. JF, JZ, SS and GA performed the tissue microdissections. SS and GA carried out the RNAscope wet lab assays. SS and GA did the confocal image capturing and analysis. SS wrote the manuscript. RSB, SS, PB and GA proofread the manuscript. SS, RSB and PB obtained the funding for research. All authors read and approved the final manuscript.

## Acknowledgements

The authors acknowledge to the NDSU-Advance Imaging Lab (AIM) for equipment support.

